# mTOR-RhoA signalling impairments in direct striatal projection neurons induce altered behaviours and striatal physiology in mice

**DOI:** 10.1101/858712

**Authors:** Daniel Rial, Emma Puighermanal, Emmanuel Valjent, Serge Schiffmann, Alban de Kerchove d’Exaerde

## Abstract

As an integrator of molecular pathways, mTOR has been associated with diseases including neurodevelopmental, psychiatric and neurodegenerative disorders as autism, schizophrenia, and Huntington’s disease. An important brain area involved in all these diseases is the striatum. However, the mechanisms behind how mTOR is involved in striatal physiology and its relative role in distinct neuronal populations in these striatal-related diseases still remain to be clarified.

Taking advantage of the D_1_-mTOR KO mice (males), we combined behavioural, biochemical, electrophysiological and morphological analysis aiming to untangle the role of mTOR in direct pathway striatal projection neurons (dMSNs) and how this would impact on striatal physiology.

Our results indicate deep behavioural changes in absence of mTOR in dMSNs such as decreased spontaneous locomotion, impaired social interaction and repetitive behaviour. These were accompanied by a Kv1.1-induced increase in the fast phase of afterhyperpolarization and decreased distal spines density that were mechanistically independent of protein synthesis but dependent of RhoA activity.

These results identify mTOR RhoA signaling as an important regulator of striatal functions through an intricate mechanism involving RhoA and culminating in Kv1.1 overfunction, which could be targeted to treat striatal-related mTORopathies.

## Introduction

The mammalian target of rapamycin (mTOR) is a protein kinase that acts as a central sensor of cell growth and metabolism, and regulates neuronal morphogenesis through the precise control of protein synthesis (1–3). mTOR serves as a core component of two different complexes, mTORC1 (sensitive to rapamycin) and mTORC2, which control distinct cellular processes. The stimulation of several G-protein coupled receptors such as D_1_R –but not D_2_R– can trigger mTORC1 activation (4). Upon activation, mTORC1 promotes protein synthesis by phosphorylating translation regulators, including p70S6K and 4EBP1 (5, 6). In addition to its control in mRNA translation, mTORC1 has been described as a postsynaptic voltage sensor, controlling the excitability of neurons (7). In addition, RhoA, which recently has been mechanistically connected to autistic-like behaviour (with marked modifications on locomotion and social skills) in mice after the deletion of the TAOK2 gene (8) and an important mediator of protein recycling in mammalian cells, is connected to the activity of the mTORC1 pathway (9). The second complex, mTORC2, has a lower sensitivity to rapamycin and its activation results in increased phosphorylation of Akt at the serine 473 site (10). It controls cell survival and the dynamics of actin cytoskeleton, which among others have been implicated in the structural plasticity of dendritic spines, including those in MSNs (11–13).

In general, monogenic disorders associated with the mTOR signalling pathway, or “mTORopathies”, match the physiological complexity and heterogeneity of the mTOR function. These are commonly divided into *i*) neurodevelopmental, such as some forms of autism (14, 15) and epileptic encephalopathy (16), *ii*) psychiatric, including schizophrenia (17) and major depressive disorder (18) and *iii*) neurodegenerative disorders, such as Alzheimer’s, Parkinson’s and Huntington’s disease (19–21). Many of these disorders are known to have striatum malfunction as a major factor of their development (22–26).

The striatum is the gateway to the basal ganglia and receives converging sensory, emotional and cognitive information via cortical and thalamic inputs. These glutamatergic synapses act on two major striatal populations of medium spiny neurons: the striatonigral cells (dMSNs), which give rise to the <u>d</u>irect pathway that mainly promotes motor output, and the striatopallidal neurons (iMSNs; dopamine D_2_R/A_2a_R-expressing neurons), which belong to the indirect pathway, conventionally classified as inhibiting motor output.

Given that mTOR controls the signalling associated with striatal function here we deleted mTOR specifically from D_1_R-expressing MSNs (dMSNs) and examined animals’ behaviour, as well as cell physiological and morphological properties. We found that the deletion of mTOR in dMSNs induces behavioural impairments associated to selective Kv1.1-channel mediated changes in dMSNs electrophysiological properties, the latter also noted in rapamycin-treated slices. Those observations also coincided with decreased density of spines along distal dendrites of dMSNs. These changes were independent of protein synthesis, but sensitive to the RhoA-dependent control of K^+^ channels recycling.

## Methods and Materials

### Animals

The mTOR floxed (mTOR^LoxP/LoxP^) mice used in the present study were described before(27). The mTOR^**Δ**/LoxP^ were obtained by crossing mTOR^LoxP/LoxP^ mice with heterozygous PGK-Cre^+/-^ mice(28) and the resulting PGK-Cre^−/-^ mTOR^**Δ**/wt^ mice were crossed with mTOR^LoxP/LoxP^ mice. The Cre-mediated deletion of mTOR in dMSNs was obtained by crossing Drd1-Cre heterozygous mice(29) and mTOR^**Δ**/LoxP^ mice. mTOR D_1_-driven knockouts (Drd1-Cre^+/-^ mTOR^**Δ**/LoxP^) and control littermates (mTOR^**Δ**/LoxP^) were obtained by crossing Drd1-Cre^+/-^ mTOR^**Δ**/LoxP^ mice with mTOR^LoxP/LoxP^ mice.

For electrophysiological and consequent 3D reconstruction experiments, to label dMSNs we crossed Drd1-Cre^+/-^ mTOR^**Δ** /LoxP^ with Ai9, Rosa26dTomato mice (30) allowing the expression of td-Tomato in neurons expressing Cre recombinase only. Drd1-Cre^+/-^ mTOR^**Δ**/LoxP^ Rosa26dTomato^+/-^ were used as D_1_-mTOR KO and control mice are Drd1-Cre^+/-^ mTOR^**Δ** /wt^ Rosa26dTomato^+/-^. All mice were of C57BL6J genetic background. All procedures were performed in accordance with the guidelines provided by the Institutional Animal Care Committee and were approved by the local ethics committees (ULB Pôle Santé – Protocol 551N). 2 to 4 mice were housed together, with *ad libitum* access to food and tap water, temperature: 21°C, humidity: 50%, light/dark cycle 12h/12h. Only males were used for the experiments.

### Western blot analysis

The striata of mice were rapidly dissected as described (31) on an ice-cooled disc, sonicated in 300 μl 10% sodium dodecyl sulfate (SDS) and boiled at 100° C for 10 min. In each experiment, samples from all animal groups were processed in parallel to minimize inter-assay variations. Protein quantification and western blots were performed as described (32, 33). All the information about the antibodies, suppliers, dilutions and references are summarized in the Supplementary Table 1.

### Puromycin incorporation in whole striatal lysates

Puromycin incorporation method was performed as described (32, 34). Briefly, the striata of mice were rapidly dissected and homogenized using 20 up-and-down strokes of a pre-chilled glass homogenizer with 800 μl of polysomal buffer containing 50 mM Tris pH = 7.8, 240 mM KCl, 10 mM MgCl_2_, 250 mM D-sucrose, 2% Triton X-100, 20 μl/ml emetine, 5 mM DTT, 100 U/ml RNasin (Promega) and protease inhibitor cocktail (Roche). Samples were centrifuged for 5 min at 16,100 × g at 4°C and supernatant was incubated with 100 μg/ml of puromycin for 10 min at 4°C and then boiled for 10 min at 100°C.

### Spontaneous locomotor activity

Horizontal and vertical activity was measured for a period of 120 minutes in a circular corridor (Imetronic, Pessac, France) as previously described (35). Counts for horizontal activity were incremented by consecutive interruption of two adjacent beams placed at a height of 1 cm per 90° sector of the corridor and counts for vertical activity (rearings) as interruption of beams placed at a height of 7.5 cm along the corridor (mice stretching upwards).

### Accelerating rotarod

Mice were placed on a rotarod apparatus (Columbus Instruments, USA) accelerating from 4 to 40 rpm in 5 min. Trials began by placing the mouse on the rod and beginning rotation. Each trial ended when the mouse fell off the rod, and latency was recorded. Mice were tested for four trials a day (1-min inter-trial interval) for 3 consecutive days (36).

### Elevated plus maze

Anxiety-like behaviour was measured as previously described (37). The elevated plus maze test was performed in a black Plexiglas apparatus with four arms (29 cm long, 5 cm wide) set in cross from a neutral central square (5 cm × 5 cm) elevated 40 cm above the floor.

### Marble burying test

Repetitive behaviour was assessed by marble burying activity. Mice were placed in a clean cage (46 × 20 × 14 cm) with 4 cm sawdust bedding overlayed by 20 glass marbles (15 mm diameter) equidistant in a 4 × 5 arrangement. Mice were allowed to explore the cage for 20 min and the number of marbles buried (>50% of the marble covered by the bedding) was counted.

### Three-chamber social approach

A three-chamber arena was used to assess sociability, preference for social novelty, and social memory as previously (38). The discrimination index for sociability was calculated as (time exploring Stranger#1 – time exploring empty wire cage) / (total exploration time) * 100 and social novelty as (time exploring Stranger#2 – time exploring Stranger#1) / (total exploration time) * 100.

### Brain slice preparation and Patch-clamping

All the procedures were similar to those described before(39). 3 months-old animals were used for these experiments. Coronal striatal slices were left in the chamber at 34°C to recover for at least 1 h before recording. In the set of experiments involving rapamycin (200 nM, Sigma-Aldrich, Germany) slices were let to recover in rapamycin-infused aCSF or DMSO (control group) for 75 min as described before(40). In the case of anisomycin (50 μM, Sigma-Aldrich, Germany) protocol, rapamycin was added to aCSF 30 min after anisomycin and 75 additional min were counted. C3 exoenzyme from *Clostridium Botulinum* (0.5 µg/ml, Cytoskeleton Inc., #CT04) was infused in aCSF for 4 h before any combined treatment following instructions of the manufacturer.

After the recovery period, slices were placed in a submersion-type chamber at room temperature with 2 ml/min rate of aCSF saturated with 95% O_2_ and 5% CO_2_. Biocytin (0.5%) was added aiming 3D reconstruction of patched-neurons. Access resistance, membrane resistance and membrane capacitance were extracted from voltage-clamp recordings by analysing current relaxation induced by a 10 mV step change from a holding potential of −80 mV as described previously (41). Intrinsic excitability was investigated in current-clamp by setting the resting membrane potential at −80 mV and injecting 1 s depolarizing steps (from 0 to 250 pA in 10 pA increments). The fast phase of afterhyperpolarisation potentials (fAHP) was calculated as described (42). Tetraethylammonium chloride (TEA, 50 μM, Sigma-Aldrich, Germany) and Dendrotoxin-K (30 nM, Sigma-Aldrich, Germany), stromatoxin (Alomone, Jerusalem, Israel) or Phrixotoxin-2 (Alomone, Jerusalem, Israel) were infused into the aCSF solution and recordings performed 10 minutes after. The concentration of TEA was based on a previous work of Hadley and colleagues (43). All recordings were analyzed using the IgorPro® 6.3 software (Wavemetrics, PO, USA).

### Immunohistochemistry and 3D-reconstruction

We used a slightly modified version of the protocol used before in our laboratory (44). Slices were incubated overnight at 4°C in paraformaldehyde solution (4% in 0.01 mM PBS). After fixation, slices were washed 3 times in PBS and incubated in PBS-0.1% TritonX-100 (Sigma-Aldrich, Germany) for 2 h under gently shaking. Slices were then transferred in solution containing streptavidin-coupled Alexa Fluor 488 (1/200 in PBS-0.1% TritonX-100 solution, Jackson ImmunoResearch, BA, USA) or NL557 (1/3000 in PBS-0.1% TritonX-100 solution, R&D Systems, Germany) for 1 h under shaking.

After the acquisition, the image was deconvoluted (Huygens-II; Scientific Volume Imaging, Netherlands) and the 3D image rendered. Reconstruction of the dendritic arborization of the neurons was done by the computer-aided filament tracing function in the 3D image analysis software Imaris (Bitplane, Ireland). A complementary manual tracing strategy adjusted the automatic reconstruction. Total dendritic arborization reconstruction allowed computation of dendrite branching, dendrite segment length analysis, total spine number and spine density in proximal (<50 μm distance from the soma) or distal dendritic domains (>50 μm distance from the soma).

### Statistical analysis

All data are expressed as mean ± standard error of the mean and the choice of the statistical tests were guided by the Shapiro–Wilk’s W normality test. If parametric, the Student’s t-test or the analysis of variance (ANOVA) followed by the Tukey test was performed. Non-parametric tests such as the Mann-Whitney and Kruskal-Wallis were followed by the Dunn’s test. All tests were performed using the Graph Pad Prism 6 software package (Graph Pad, San Diego, CA, USA). The accepted level of significance for the tests was P ≤ 0.05. All descriptive statistical results are summarized in the Supplementary Table 2.

## Results

### Deletion of mTOR in dMSNs results in decreased activity of downstream targets of mTORC1 and modifies the behaviour of mice

mTOR as a complex hub of molecular interactions is capable of modulating targets in control of protein synthesis, membrane dynamics and cell excitability (6). We therefore evaluated the downstream molecular consequences of the inactivation of mTOR in dMSNs by using western blot analysis. As expected, a significant reduction of mTOR expression was observed in the striatum of D_1_-mTOR KO (Drd1 Cre^+/-^ mTOR^**Δ** /LoxP^) compared to control littermates (mTOR^**Δ**^ /^LoxP^) (Figure 1A,F). Additionally, decreased phosphorylation levels of two major downstream effectors of mTORC1 namely p70S6K (phosphorylated at threonine 389) (Figure 1B,F) and 4EBP1 (phosphorylated at threonine 37/46) (Figure 1C,F) were observed in the striatum of D_1_-mTOR KO mice. In contrast, no difference was observed in the phosphorylation state of Akt at serine 473 suggesting that mTORC2 activity was not altered in the striatum (Figure 1D,F). Moreover, puromycin-based assay revealed that global protein synthesis was not altered in the striatum of D_1_-mTOR KO mice (Figure 1E).

**Figure 1.**
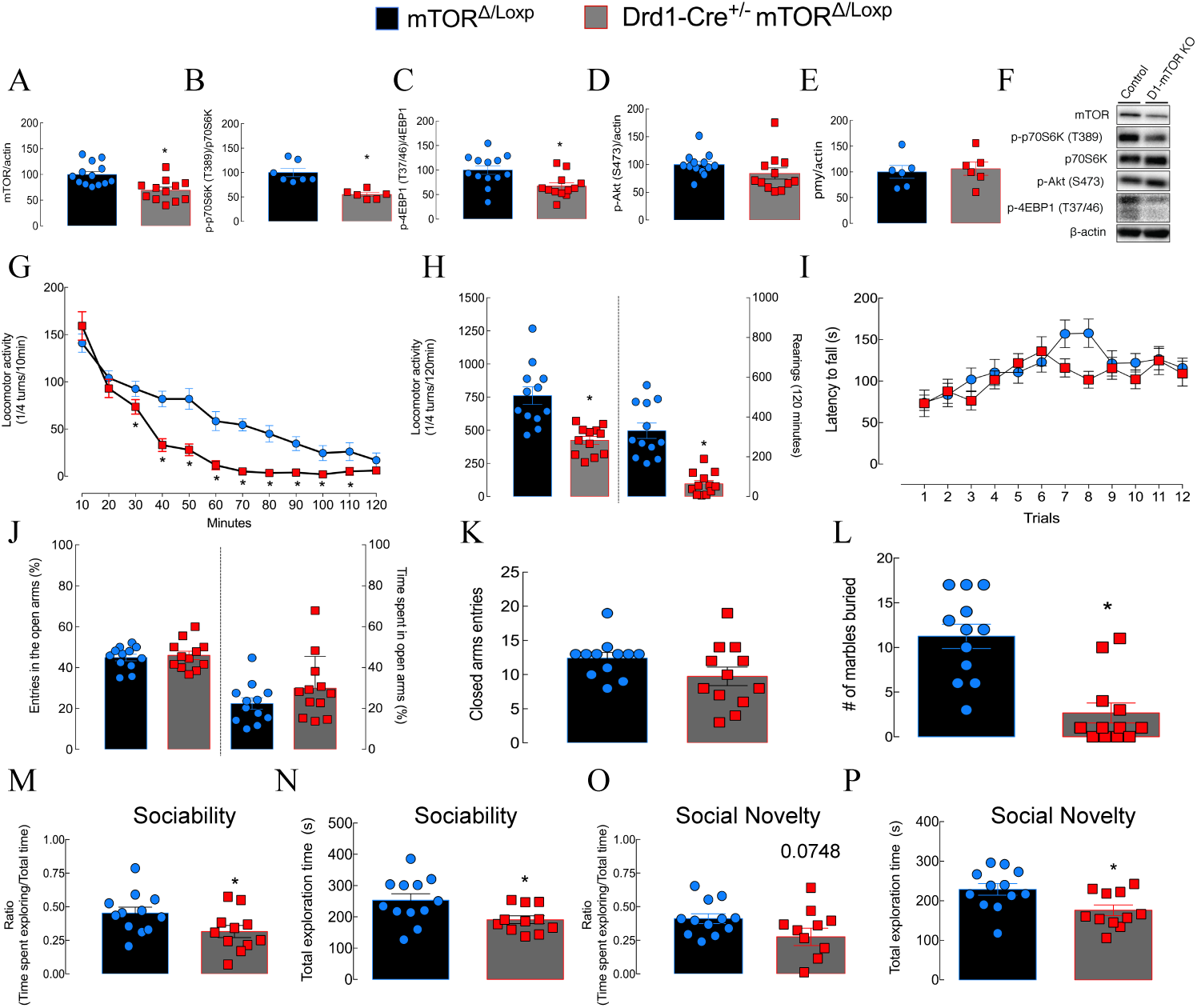
Deletion of mTOR decreases spontaneous locomotor activity, modifies repetitive and social behaviour in mice. The reduction in the expression of mTOR in the total striatum of dMSNs mTOR-KO mice is shown in (A). After the D_1_-driven deletion of mTOR, lower expression of mTOR-partner proteins such as p-p70S6K (phosphorylation site T389) (B) and p-4EBP1 (phosphorylation site T37/46) (C) was observed. The phosphorylation pattern of Akt (phosphorylation site S473) is depicted in (D). (E) Similar levels of puromycin incorporation in both groups. (F) Representative western blots. Decreased horizontal and vertical spontaneous locomotor activity in mTOR-KO mice when compared to control littermates in the circular corridor test (G and H). The decrease in the locomotor activity is not related with deficits of motor coordination and motor learning as can be observed by our rota-rod results (I) or anxiety-like behaviour (J and K). Repetitive behaviour evaluated by the marble-burying test revealed that mTOR-KO mice buried less marbles than control mice (L). The deletion of mTOR in dMSNs induces the reduction of social interaction with conspecific mice (M) and (N). Social novelty (when a second conspecific mouse is introduced) is partially impaired following the mTOR deletion in dMSNs (O) and (P). * p<0.05.

Since it has been previously described that the manipulation of upstream and downstream mTOR regulators affect locomotor activity (45), we evaluated the voluntary locomotor activity of D_1_-mTOR KO and control littermates. Our results revealed that the selective deletion of mTOR in dMSNs significantly impaired spontaneous locomotor behaviour (Figure 1G). However, to note is the difference in the locomotor curves over time indicating similar locomotor activity during the first 20 minutes of analysis but exponential decay kinetics in mTOR-KO mice. Figure 1H shows the quantification of the total ambulation during a 120 minutes session corroborating the decreased locomotion in D_1_-mTOR KO mice and reveals a decrease in vertical activity in the same group when compared to control mice. This reduced locomotor activity was not associated with increased anxiety levels since no alterations were observed in the elevated-plus maze (Figures 1J and K). Finally, animals were submitted to the marble burying task and we found that D_1_-mTOR KO buried less marbles than control mice, indicating altered repetitive behaviour (Figure 1L). Interestingly, D_1_-mTOR KO displayed similar motor performance as littermates on the rotarod suggesting that not all repetitive motor routines were affected (Figure 1I).

Next, we decided to evaluate the sociability and social novelty behaviour in both groups. Results gathered from the three-chamber social apparatus revealed decreased sociability in D_1_-mTOR KO mice in comparison to control littermates (Figure 1M) accompanied by a reduction of the total exploration time (Figure 1N). Social novelty might have been also impaired in D_1_-mTOR KO given that there was a trend in the time exploring/total time ratio (Figure 1O) and a significant decrease in the total exploration time (Figure 1P) was observed.

Together, our results indicate that the deletion of mTOR in dMSNs reduces voluntary locomotion accompanied by disturbances in repetitive and impaired social behaviour without modifications in the anxiety profile of mice.

### Inactivation of voltage-gated potassium channels 1.1 (Kv1.1) rescues the neurophysiological abnormalities induced by genetic or pharmacological mTOR inhibition

The striatal encoding of locomotor activity is causally connected to the excitable properties and activity of dMSNs and iMSNs(46). The control of excitability exerted by mTOR relates to the intimate control of voltage-gated potassium channels (Kvs) (47). The Kv1 family are important components controlling the somatodendritic excitability in sensory neurons (48) and MSNs (49). Importantly, mTOR has been shown to dampen excitability by increasing Kv1.1 channel function in hippocampal neurons (7, 40). Hence, we hypothesized that some of the behavioural alterations observed in D_1_-mTOR KO mice are caused by changes in electrophysiological properties. We therefore carried out the electrophysiological characterization of dMSNs in presence and absence of mTOR. First, we observed that resting membrane potentials (RMP) were similar between groups (Figure 2A). Interestingly, a 40% reduction in capacitance (proportional to the cell size) was observed in dMSNs lacking mTOR as compared to control dMSNs (Figure 2B). Our electrophysiological data also identified an increase in the amplitude of the fast phase of afterhyperpolarization (fAHP) (Figure 2C). To probe for a direct and acute control by mTOR of these electrophysiological changes, we employed an *in vitro* protocol where slices from both genotypes were submitted to an incubation with the mTOR inhibitor rapamycin prior the recordings, as described before (40). We first confirmed the lack of modification in the RMP in the presence of rapamycin or DMSO (Figure 2D). Rapamycin did not affect the cell capacitance either in control or mTOR-KO dMSNs (Figure 2E). Acute *in vitro* exposure to rapamycin in control mice phenocopied the higher amplitude of the fAHP of dMSNs in D_1_-mTOR KO. In contrast, rapamycin exposure had no effect on the enhanced fAHPs of dMSNs in D_1_-mTOR KO (Figure 2F). We also computed other active properties of dMSNs such as action potential (AP) threshold, amplitude, duration, depolarization and hyperpolarization rate and input-output curve (data not shown) without any differences between groups. To test if this increase in the amplitude of fAHP was due to a higher density/function of potassium channels in D_1_-mTOR KO and rapamycin-treated control dMSNs we first used low concentrations of TEA (a non-selective Kv channels blocker) and Dendrotoxin-K (a highly selective Kv1.1 channel blocker). The concentration of 30 nM of Dendrotoxin-K was selected based on the curve showed in the Figure 2G aiming a concentration with minimal effects in control dMSNs but that would exhibit effects in a situation where increased density or function of Kv1.1 is present. Figure 2H shows that in control D_1_ Cre^+/-^ mTOR^Δ/wt^ Rosa26dTomato^+/-^ dMSNs, the rapamycin-induced increase in the amplitude of fAHP is re-normalized by the non-selective blockade of Kv-channels and the selective blockade of Kv1.1 channels. The same analysis in D_1_ Cre^+/-^ mTOR^Δ/LoxP^ Rosa26dTomato^+/-^ KO cells indicated that the higher amplitude of the fAHP is promptly reduced by TEA and Dendrotoxin-K to similar level than in control dMSNs in basal conditions. The effects were similar in DMSO- and rapamycin-treated mTOR-KO dMSNs confirming again the absence of additive effects. To further check for additional channels that could be involved in the fAHP changes, we used the *in vitro* rapamycin protocol in control dMSNs and recorded in the presence of Stromatoxin (a selective Kv4.2 blocker) and Phrixotoxin (a blocker of Kv4.1 and Kv4.3). After building the respective inhibition curves to set the concentrations of both drugs to be used (respectively Figures 2I and J) using the same rationale as for the Dendrotoxin-K, our electrophysiological analysis indicated that Kv4.1, 4.2 and 4.3 channels were not involved in the mTOR-induced increase in the fAHP amplitude (Figure 2K). The electrophysiological analysis indicates deep modifications in mTOR-KO dMSNs and rapamycin-treated dMSNs with alterations in the capacitance and fAHP that is Kv1.1-related.

**Figure 2.**
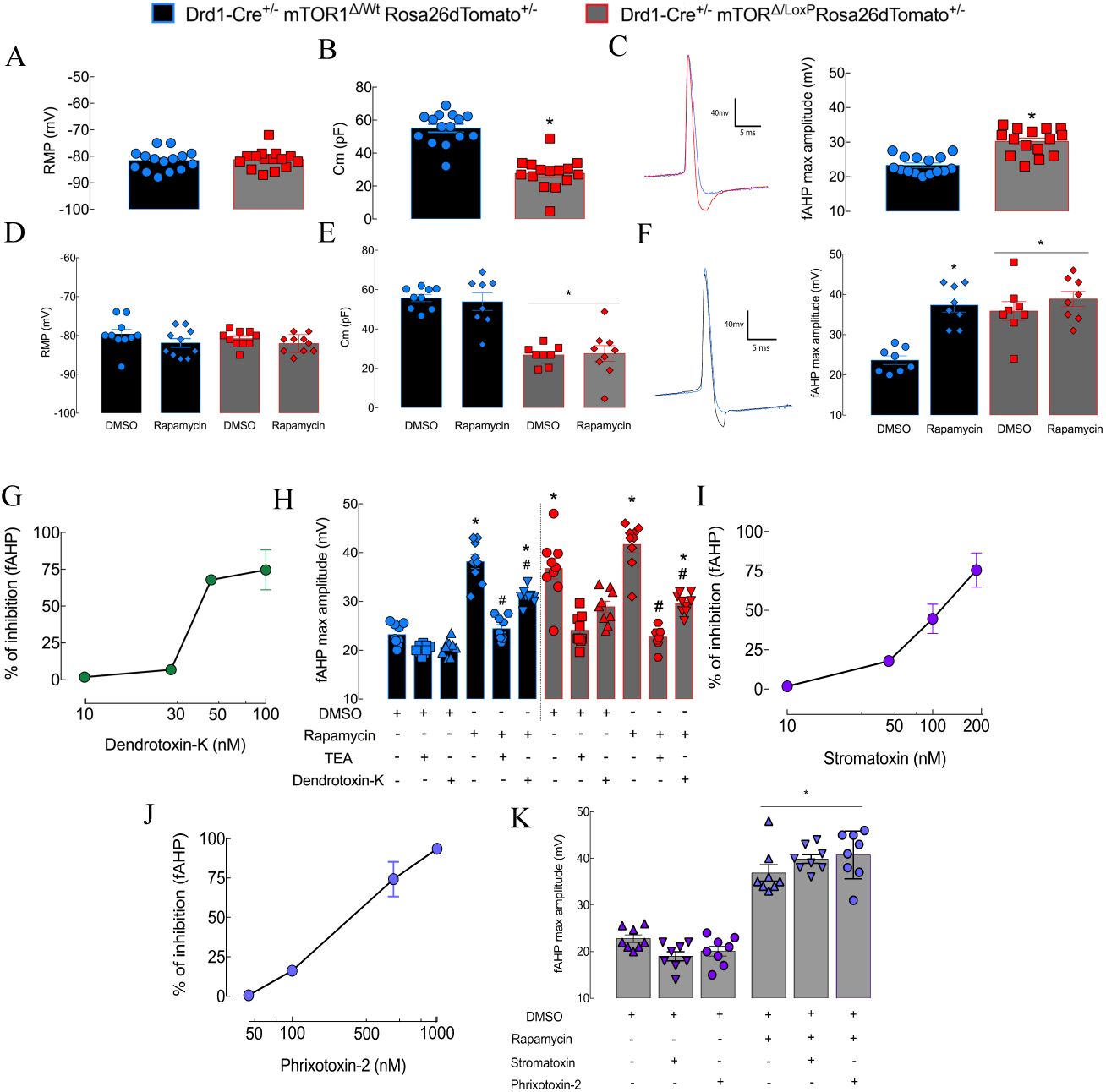
Kv1.1 mediates the electrophysiological modifications in mTOR-KO dMSNs. The deletion of mTOR in dMSNs did not modify the RMP of these neurons (A) however it decreased cell capacitance (B) and increased the amplitude of fAHP of dMSNs from mTOR-KO mice (red trace) in comparison to control dMSNs (blue trace) (C). The pharmacological blockade of mTOR did not produce any effects on the RMP (D) or additive effects on the capacitance of both genotypes (E) but promptly increased the amplitude of the fAHP (rapamycin: black trace and DMSO: blue trace) similar to the genetic manipulation (F). After the selection of the appropriate concentration of TEA and dendrotoxin-K (G), panel H shows that, the non-selective blockade of Kvs (TEA) completely renormalized the mTOR-mediated increase in the fAHP, effect that is partially recapitulated by the selective blockade of Kv1.1 (dendrotoxin-k). A similar approach but aiming the Kv4.2 (J) and Kv4.1 and 4.3 showed that these channels are not involved in the mTOR-mediated increase of fAHP (K) (Stromatoxin 50 nM and Phrixotoxin-2 100 nM). *p<0.05 when compared with the DMSO control group. # p<0.05 when compared with the respective genotype rapamycin exposed cells.

### Reduced dendritic arborisation and spine density in dMSNs lacking mTOR activity

Cytoskeleton reorganization is controlled by mTOR (50) and evidence indicates that mTOR-dependent actin polymerization through RhoA mechanisms controls spine formation and maturation (51). Moreover, the increased amplitude of the fAHP coupled with the lack of changes in somatic excitability could be accounted by morphological changes in distal dendritic branches (52). We therefore performed 3D reconstruction of the recorded dMSNs (Figure 3A-C). Computation of the dendritic arborization and classification in different branch levels (Figure 3D) show that mTOR-KO dMSNs are less ramified when compared to the control counterparts and that rapamycin has no impact on this parameter. However, both the genetic deletion of mTOR in dMSNs and the pharmacological blockade of mTOR by rapamycin in control animals decreased the total number of spines in dMSNs (Figure 3E). Rapamycin did not decrease further the number of spines in mTOR-KO dMSNs showing the absence of additive effects. A detailed evaluation of the spine density, close (< 50 μm) to the soma (Figure 3F) or far from the soma (> 50 μm) (Figure 3G), revealed similar values for all groups for proximal spines but a significant lower distal spine density in rapamycin-treated control dMSNs and mTOR-KO dMSNs. The morphological data indicates selective distal dendritic changes in dMSNs after the genetic and pharmacological blockade of mTOR in agreement with the fAHP changes.

**Figure 3.**
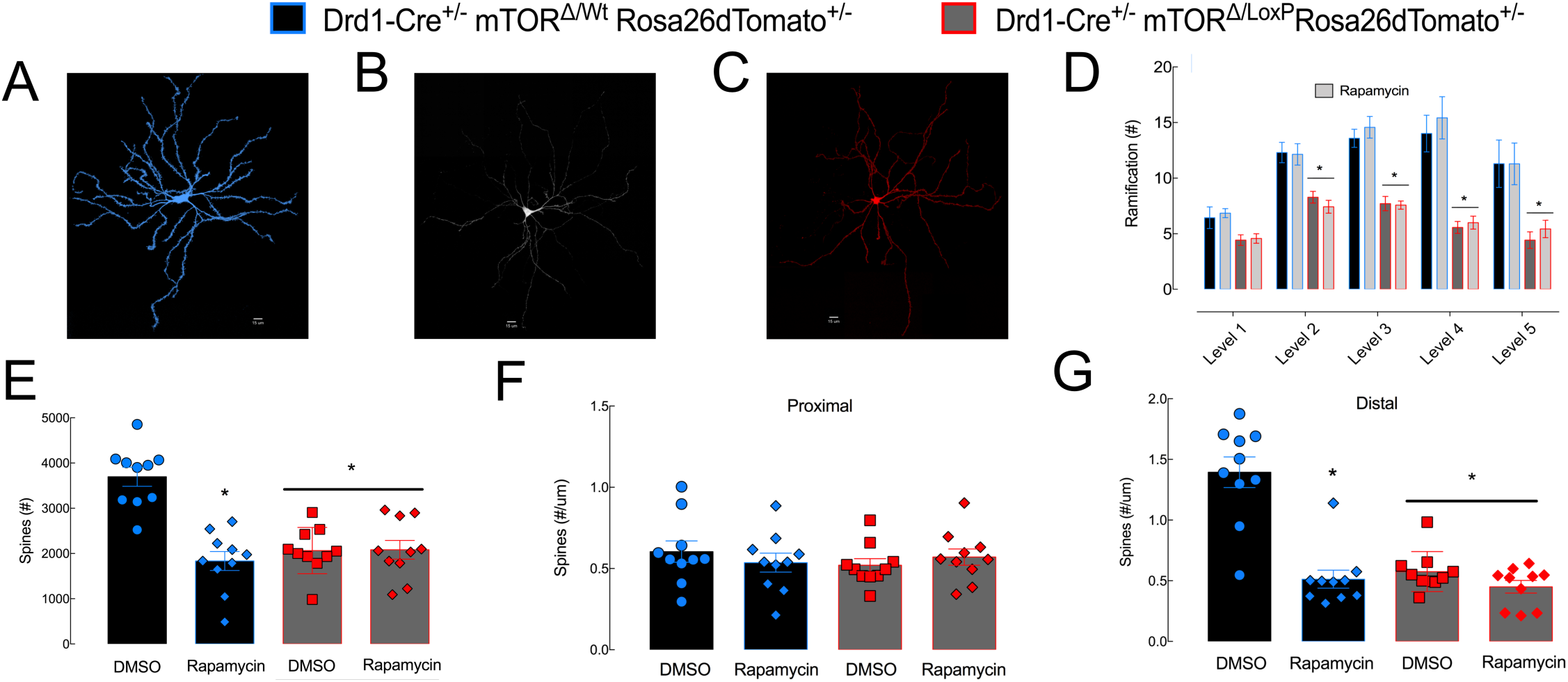
Selective morphological characteristics are shaped by mTOR in dMSNs. Representative 3D reconstructions of dMSNs of control (A), rapamycin (B) and D_1_-mTOR KO (C) conditions are shown. Decreased ramification of dMSNs after the genetic deletion of mTOR but no effect of rapamycin on this parameter was noted (D). Total number of spines in dMSNs is decreased after the exposure to rapamycin or the genetic deletion of mTOR (E). Proximal (<50 μm from the soma) spine density is not affected by rapamycin or the mTOR deletion in dMSNs (F) but distal spine density (>50 μm) is impaired by both conditions (G). *p<0.05 when compared with DMSO control group.

### Electrophysiological and morphological changes induced by the lack of mTOR activity in dMSNs are independent of protein synthesis

We next investigated whether electrophysiological and morphological changes in dMSNs provoked by the pharmacological mTOR inhibition are dependent of protein synthesis. Our data show that the *in vitro* exposure to anisomycin (a protein synthesis inhibitor) prior to the recordings did not affect the rapamycin-induced enhancement of Dendrotoxin-sensitive fAHPs. Similar RMP values were found in all groups (Figure 4A) and a decreased capacitance in dMSNs from Drd1-Cre^+/-^ mTOR^**Δ**/LoxP^ Rosa26dTomato^+/-^ (Figure 4B) was still present when compared to dMSNs from control littermates. Remarkably, the increase in the amplitude of the fAHP is maintained in the presence of anisomycin after pharmacological blockade or genetic deletion of mTOR (Figure 4C). Morphological data showed that the total number of spines as well as spine density far from the soma remained reduced after the pharmacological blockade and the genetic deletion of mTOR in dMSNs regardless of the protein synthesis inhibition (Figures 4D and F respectively). On the other hand, there is still no impact on the proximal spine density (Figure 4E). The lack of protein synthesis involvement in these alterations suggests alternative mechanisms such as channel recycling.

**Figure 4.**
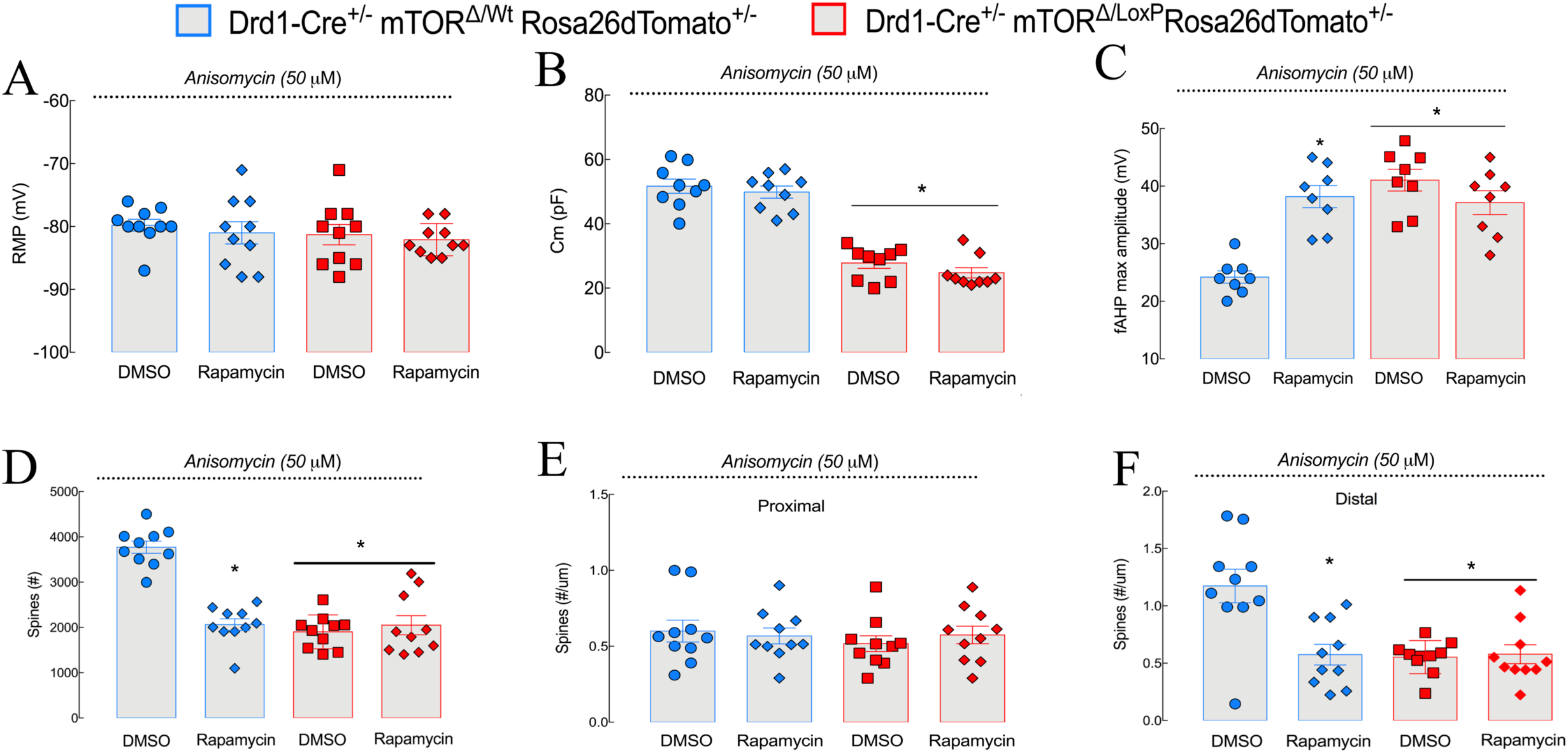
mTOR-induced electrophysiological and morphological modifications are protein-synthesis independent. The exposure to anisomycin (a protein synthesis inhibitor) did not change the RMP of control and mTOR KO dMSNs or changed the profile of the response for the capacitance (B) or the fAHP amplitude (C). Similarly, no changes in the proximal spine density (D) were observed while the decreased in the total number of spines (E) and distal spine density (F) are maintained regardless of the exposure to anisomycin. * p<0.05 when compared with the DMSO control group.

### Inhibition of RhoA-mediated channel recycling prevents rapamycin-induced electrophysiological and morphological changes

We then addressed if the observed electrophysiological and morphological alterations are due to the disturbance of RhoA-mediated membrane recycling mechanisms by incubating slices with Clostridium botulinum C3 transferase exoenzyme (C3 exo), a RhoA inhibitor. There are no effects of C3 exo on the RMP (Figure 5A), or on the dMSNs capacitance (Figure 5B), which remained reduced in D_1_ Cre^+/-^ mTOR^Δ/LoxP^ Rosa26dTomato^+/-^ mice. On the other hand, exposure to C3 exo prevented the rapamycin-induced increase in the fAHP amplitude in dMSNs from control D_1_ Cre^+/-^ mTOR^Δ/wt^Rosa26dTomato^+/-^ mice but did not modify the increased fAHP amplitude of mTOR-KO dMSNs (Figure 5C). 3D reconstruction of dMSNs revealed that exposure to C3 exo prevented the rapamycin-induced reduction of the total spine number and distal spine density (Figures 5D and F) in control dMSNs without particular changes in the reduced levels in mTOR-KO dMSNs. The proximal spine density was not impacted by C3 exo (Figure 5E). Taken together, these results show that the rapamycin effects on fAHP and dMSNs morphology rely on RhoA-dependent mechanisms.

**Figure 5.**
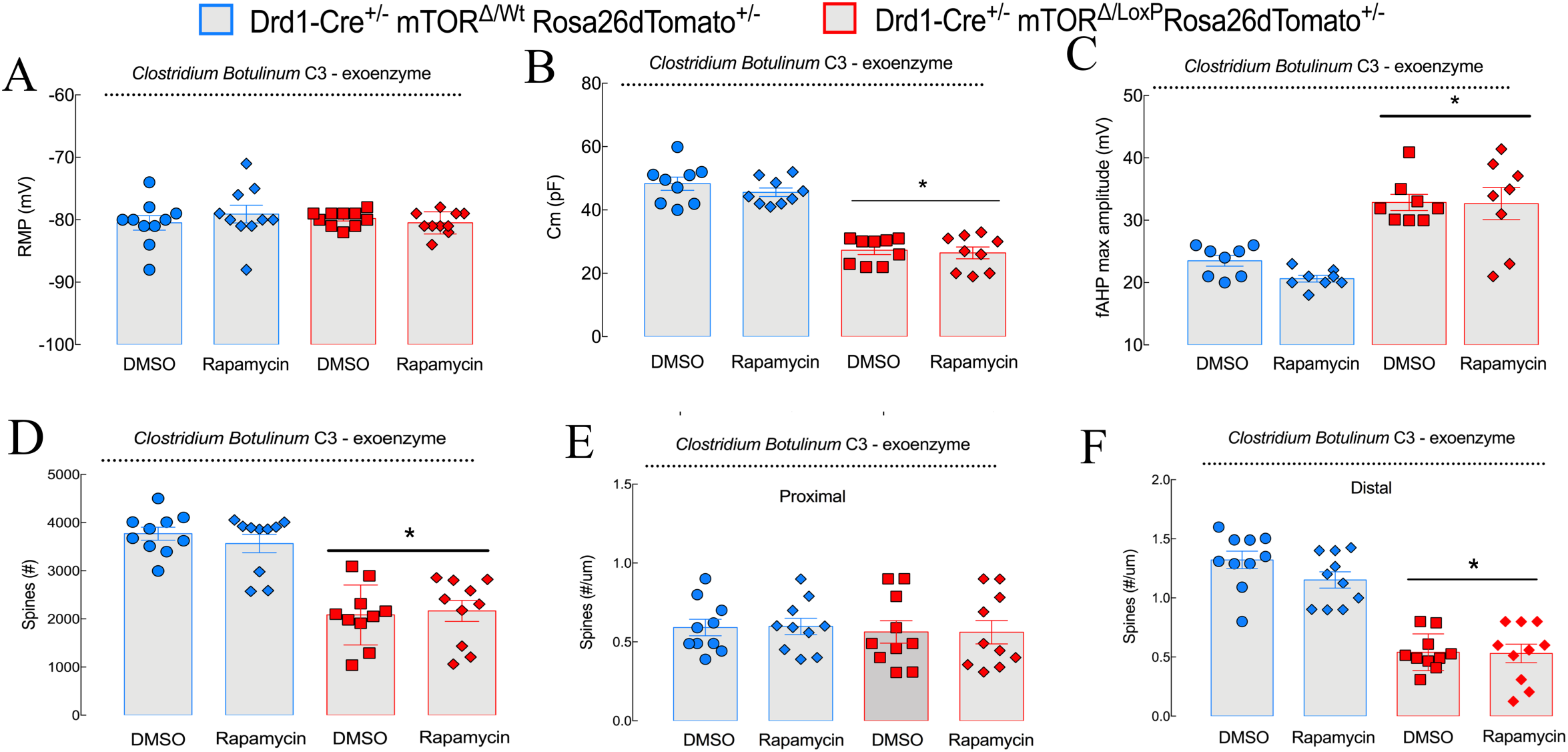
RhoA inhibition prevents the rapamycin-induced electrophysiological and morphological modifications. The exposure to the C3 exoenzyme (a RhoA inhibitor) was not sufficient to induce changes in the dMSNs RMP (A) interfere with the capacitance (B). However, it prevented the rapamycin-induced increase of fAHP amplitude (C) without interfering with the same parameter in mTOR KO dMSNs. The same pattern of response was seen in the total number of spines (D) and distal spine density (F) without inducing changes in the proximal spine density (E). *p<0.05 when compared to the DMSO control group.

## Discussion

In this study, we determined the role of mTOR in dMSNs by assessing how the genetic deletion of mTOR from dMSNs affects striatal-related behaviours and their neurophysiology. Our behavioural data indicate that the inactivation of mTOR in dMSNs disrupts several aspects of striatal circuitry physiology. Using a cell-type-specific approach, we unveiled that mTOR controls, through a protein synthesis-independent mechanism, the membrane density of Kv1.1 using RhoA as substrate. Moreover, the genetic deletion of mTOR in dMSNs induces modifications of the dendritic complexity and distal spine density, resulting in behavioural changes compatible with disrupted striatal function. Additionally, we found that short-term exposure of dMSNs from control mice to rapamycin, a specific mTOR inhibitor (53, 54), replicates electrophysiological and morphological features found in neurons genetically inactivated for mTOR.

Our first aim was to determine the functional consequences of the selective mTOR inactivation in dMSNs. Our findings that D_1_R-mTOR KO mice exhibit impaired social and repetitive behaviours are reminiscent of phenotypes observed in several forms of idiopathic autism disorders in which aberrant activity of mTOR signalling has been reported (55). However, the reduced repetitive behaviours and voluntary locomotion indicate that the phenotype displayed by the D_1_R-mTOR KO mice is also intricately linked to motor dysfunctions. Sutton and Caron have previously shown that the selective deletion of mTOR or Raptor (a partner of mTORC1) in D_1_R MSNs neurons decreases locomotion in mice (56). Another mTORC1 partner, Rhes, has also been implicated on the striatal control of locomotion (57) reinforcing the control of locomotor activity by mTOR and downstream partners. However, the acute administration of rapamycin does not impact the locomotion of mice (58) suggesting that modifications after the disruption of mTOR signalling are timely dependent (development included) or specific to certain brain areas or neuronal types. Alternatively, the lack of cell-type selectivity (on D_1_ and D_2_ neurons) after rapamycin administration might be compensated by secondary mechanisms stabilizing the locomotor function, which cannot be reached by the D_1_ selective genetic deletion of mTOR.

The modifications in the activation of core components of the translational machinery such as p-4EBP1 (59) and p-p70S6K (60) could induce a set of electrophysiological alterations since mTOR controls protein synthesis and cell excitability. However, global changes in protein synthesis are unlikely in our animals since the puromycin incorporation assay indicated similar values for both groups. Kv1.1 function is known to be controlled by mTOR in a protein-synthesis independent manner (40). Notably, the fAHP modifications induced by both the selective dMSNs genetic deletion of mTOR and the rapamycin-induced pharmacological blockade were connected to the Kv1.1 activity. The fAHP modifications found here are similar to those observed when PTEN, an upstream regulator of the Akt/mTOR pathway is mutated. However, in this report, the authors associate their results with the overexpression of calcium-activated potassium channels in the cortex (61). The close connection between low levels of mTOR signalling in the striatum and altered potassium currents has been also reported in mouse models of Huntington’s disease (HD) (62).

Although neuronal morphological parameters such as dendritic complexity, volume of the soma and spine density are governed by diverse mechanisms, both mTOR complexes are key for cytoskeletal organization and function (63, 64). Changes in mTOR signalling and spine pruning defects may represent a secondary mechanism in response to an imbalance between excitatory and inhibitory neurotransmission, identified in both Mecp2 mutant mice and Tsc1-deficient mice (both partners of mTOR) (65–67) and also linked to autism. Interestingly, the deletion of the mTOR negative regulator Tsc1 induces increased excitability and decreased dendritic complexity in dMSNs (68).

RhoA control over membrane recycling of proteins, mainly receptors and channels (69, 70), and more specifically in the regulation of the traffic of voltage-gated potassium channels has been shown before (71). Interestingly, Luykennar and colleagues described in rat cerebral arteries that the inhibition of RhoA leads to a concomitant loss of pyrimidines control of excitability, attributed on their specific case to the Kv1.2 malfunction, paired with actin disorganization (72). Still, the selective prevention of modifications exerted by C3 exo has only been observed in the rapamycin protocol but not in cells from D_1_-mTOR KO mice. Two possible explanations might account for this difference. First, developmental differences where mTOR is known to be an important factor (73), and second, as C3 exo provides prevention, this would not be observable with any approach using KO animals since the time resolution is lost.

Altogether, our results indicate that the selective deletion of mTOR in dMSNs induces deep behavioural changes accompanied by electrophysiological and morphological modifications that are dependent of an over function of Kv1.1 channels regulated by RhoA. We also reinforce the notion of mTOR as a valid target to striatal dysfunction and provide, with cell-type specificity, a mechanistic hint about the dynamic changes in MSNs electrophysiology and morphology that accompany striatal disorders.

## Acknowledgements

D.R. is a BELSPO Fellowship recipient; A.K.E is a Research Director of the FRS-FNRS and an investigator of WELBIO. FRS-FNRS (Belgium), Action de Recherche Concertée (FWB) and PAI 7/10 (Federal Science Policy) supported this study. Foundation Simone and Pierre Clerdent supported AKE. E.V. is supported by Inserm, Fondation pour la Recherche Médicale (FRM) and the French National Research Agency (ANR-EPITRACE). E.P. was a recipient of a Marie Curie Intra-European Fellowship (IEF327648), a NARSAD Young Investigator Grant from the Brain and Behavior Research Foundation and a recipient of Beatriu de Pinós fellowship (#2017BP00132) from University and Research Grants Management Agency (Government of Catalonia). We are grateful to Dr. Jochen Roeper for his comments and critical reading of the manuscript.

## Disclosure

The authors declare no conflict of interest or biomedical financial interests.

