## SupplementaryInformation for "mTOR-RhoA signalling impairments in direct striatal projection neurons induce altered behaviours and striatal physiology in mice"

**Table 1. List of primary antibodies**

| Antigen | Species | Dilution | Supplier/Catalog no./Ref |
|---|---|---|---|
| mTOR | Rabbit | 1:1,000 | Cell Signaling/#2976 |
| β-actin | Mouse | 1:40,000 | Abcam/#ab6276 |
| p-p70S6K-T389 | Rabbit | 1:1,000 | Cell Signaling/#9234 |
| p-4-EBP1-T37/46 | Rabbit | 1:500 | Cell Signaling/#2855 |
| 4-EBP1 | Rabbit | 1:500 | Cell Signaling/#9644 |
| p-Akt-S473 | Rabbit | 1:1,000 | Cell Signaling/#4060 |
| Puromycin | Mouse | 1:1,000 | David et al., 2013 [36] |

**Table 2. Descriptive statistical analysis**

| Figure | Groups (n) | Statistical analysis |
|---|---|---|
| 1A | $mTOR^{\Delta/Loxp}$ (n=13)<br>Drd1 $Cre^{+/-}$ $mTOR^{\Delta /LoxP}$ (n=12) | $t$ =3.609; df =23; p=0.0015 |
| 1B | $mTOR^{\Delta/Loxp}$ (n=7)<br>Drd1 $Cre^{+/-}$ $mTOR^{\Delta /LoxP}$ (n=6) | $t$ =4.722; df =11; p=0.0006 |
| 1C | $mTOR^{\Delta/Loxp}$ (n=13)<br>Drd1 $Cre^{+/-}$ $mTOR^{\Delta /LoxP}$ (n=12) | $t$ =2.899; df =23; p=0.0081 |
| 1D | $mTOR^{\Delta/Loxp}$ (n=13)<br>Drd1 $Cre^{+/-}$ $mTOR^{\Delta /LoxP}$ (n=12) | $t$ =1.438; df =23; p=0.1639 |
| 1E | $mTOR^{\Delta/Loxp}$ (n=6)<br>Drd1 $Cre^{+/-}$ $mTOR^{\Delta /LoxP}$ (n=6) | $t$ =0.342; df =10; p=0.7395 |
| 1G | $mTOR^{\Delta/Loxp}$ (n=12)<br>Drd1 $Cre^{+/-}$ $mTOR^{\Delta /LoxP}$ (n=12) | $F_{genotype\ (1,22)}$= 21.13, p=0.0001<br>$F_{time(11,\ 242)}$= 80.74, p<0.0001<br>$F_{interaction(11,\ 242)}$=5.071, p<0.0001 |
| 1H | $mTOR^{\Delta/Loxp}$ (n=12)<br>Drd1 $Cre^{+/-}$ $mTOR^{\Delta /LoxP}$ (n=12) | Horizontal: $t$=4.597; df=22; p<0.0001<br>Vertical: $t$=6.298; df=22; p<0.0001 |
| 1I | $mTOR^{\Delta/Loxp}$ (n=12)<br>Drd1 $Cre^{+/-}$ $mTOR^{\Delta /LoxP}$ (n=12) | $F_{interaction(11,\ 242)}$=0.2337, p=0.1545 |
| 1J | $mTOR^{\Delta/Loxp}$ (n=12)<br>Drd1 $Cre^{+/-}$ $mTOR^{\Delta /LoxP}$ (n=12) | % entries: $t$ =0.4026; df=22; p=0.6911<br>% time: $t$ =1.418; df =22; p=0.1703 |
| 1K | $mTOR^{\Delta/Loxp}$ (n=12)<br>Drd1 $Cre^{+/-}$ $mTOR^{\Delta /LoxP}$ (n=12) | $t$ =1.932; df =22; p=0.0664 |
| 1L | $mTOR^{\Delta/Loxp}$ (n=12)<br>Drd1 $Cre^{+/-}$ $mTOR^{\Delta /LoxP}$ (n=12) | $t$ =4.865; df =22; p<0.0001 |
| 1M | $mTOR^{\Delta/Loxp}$ (n=12)<br>Drd1 $Cre^{+/-}$ $mTOR^{\Delta /LoxP}$ (n=11) | $t$ =2.166; df =21; p=0.0419 |
| 1N | $mTOR^{\Delta/Loxp}$ (n=12)<br>Drd1 $Cre^{+/-}$ $mTOR^{\Delta /LoxP}$ (n=11) | $t$ =2.485; df =21; p=0.0215 |
| 1O | $mTOR^{\Delta/Loxp}$ (n=12)<br>Drd1 $Cre^{+/-}$ $mTOR^{\Delta /LoxP}$ (n=11) | $t$ =1.875; df =21; p=0.0748 |
| 1P | $mTOR^{\Delta/Loxp}$ (n=12)<br>Drd1 $Cre^{+/-}$ $mTOR^{\Delta /LoxP}$ (n=11) | $t$ =2.62; df =21; p=0.0160 |

| | | |
|---|---|---|
| 2A | Drd1-Cre$^{+/-}$ mTOR$^{\Delta/Wt}$ Rosa26dTomato$^{+/-}$ (n=15)<br>Drd1-Cre$^{+/-}$ mTOR$^{\Delta/LoxP}$ Rosa26dTomato$^{+/-}$ (n=15) | Mann-Whitney test; p=0.8293 |
| 2B | Drd1-Cre$^{+/-}$ mTOR$^{\Delta/Wt}$ Rosa26dTomato$^{+/-}$ (n=15)<br>Drd1-Cre$^{+/-}$ mTOR$^{\Delta/LoxP}$ Rosa26dTomato$^{+/-}$ (n=15) | Mann-Whitney test; p<0.0001 |
| 2C | Drd1-Cre$^{+/-}$ mTOR$^{\Delta/Wt}$ Rosa26dTomato$^{+/-}$ (n=15)<br>Drd1-Cre$^{+/-}$ mTOR$^{\Delta/LoxP}$ Rosa26dTomato$^{+/-}$ (n=15) | Mann-Whitney test; p<0.0001 |
| 2D | Drd1-Cre$^{+/-}$ mTOR$^{\Delta/Wt}$ Rosa26dTomato$^{+/-}$ (n=10)<br>Drd1-Cre$^{+/-}$ mTOR$^{\Delta/LoxP}$ Rosa26dTomato$^{+/-}$ (n=10) | Kruskal-Wallis test; p=0.3043 |
| 2E | Drd1-Cre$^{+/-}$ mTOR$^{\Delta/Wt}$ Rosa26dTomato$^{+/-}$ (n=10)<br>Drd1-Cre$^{+/-}$ mTOR$^{\Delta/LoxP}$ Rosa26dTomato$^{+/-}$ (n=10) | Kruskal-Wallis test; p=0.0015 |
| 2F | Drd1-Cre$^{+/-}$ mTOR$^{\Delta/Wt}$ Rosa26dTomato$^{+/-}$ (n=10)<br>Drd1-Cre$^{+/-}$ mTOR$^{\Delta/LoxP}$ Rosa26dTomato$^{+/-}$ (n=10) | Kruskal-Wallis test; p=0.0008 |
| 2H | Drd1-Cre$^{+/-}$ mTOR$^{\Delta/Wt}$ Rosa26dTomato$^{+/-}$ (n=10)<br>Drd1-Cre$^{+/-}$ mTOR$^{\Delta/LoxP}$ Rosa26dTomato$^{+/-}$ (n=10) | Kruskal-Wallis test; p<0.0001 |
| 2K | Drd1-Cre$^{+/-}$ mTOR$^{\Delta/Wt}$ Rosa26dTomato$^{+/-}$ (n=10)<br>Drd1-Cre$^{+/-}$ mTOR$^{\Delta/LoxP}$ Rosa26dTomato$^{+/-}$ (n=10) | Kruskal-Wallis test; p<0.0001 |
| 3D | Drd1-Cre$^{+/-}$ mTOR$^{\Delta/Wt}$ Rosa26dTomato$^{+/-}$ (n=11)<br>Drd1-Cre$^{+/-}$ mTOR$^{\Delta/LoxP}$ Rosa26dTomato$^{+/-}$ (n=11) | Kruskal-Wallis test; p=0.0174 |
| 3E | Drd1-Cre$^{+/-}$ mTOR$^{\Delta/Wt}$ Rosa26dTomato$^{+/-}$ (n=11)<br>Drd1-Cre$^{+/-}$ mTOR$^{\Delta/LoxP}$ Rosa26dTomato$^{+/-}$ (n=11) | Kruskal-Wallis test; p=0.0001 |
| 3F | Drd1-Cre$^{+/-}$ mTOR$^{\Delta/Wt}$ Rosa26dTomato$^{+/-}$ (n=11)<br>Drd1-Cre$^{+/-}$ mTOR$^{\Delta/LoxP}$ Rosa26dTomato$^{+/-}$ (n=11) | Kruskal-Wallis test; p=0.5206 |

| | | |
|---|---|---|
| 3G | Drd1-Cre$^{+/-}$ mTOR$^{\Delta/Wt}$ Rosa26dTomato$^{+/-}$ (n=11)<br>Drd1-Cre$^{+/-}$ mTOR$^{\Delta/LoxP}$ Rosa26dTomato$^{+/-}$ (n=11) | Kruskal-Wallis test; p=0.0001 |
| 4A | Drd1-Cre$^{+/-}$ mTOR$^{\Delta/Wt}$ Rosa26dTomato$^{+/-}$ (n=9)<br>Drd1-Cre$^{+/-}$ mTOR$^{\Delta/LoxP}$ Rosa26dTomato$^{+/-}$ (n=9) | Kruskal-Wallis test; p=0.4363 |
| 4B | Drd1-Cre$^{+/-}$ mTOR$^{\Delta/Wt}$ Rosa26dTomato$^{+/-}$ (n=9)<br>Drd1-Cre$^{+/-}$ mTOR$^{\Delta/LoxP}$ Rosa26dTomato$^{+/-}$ (n=9) | Kruskal-Wallis test; p<0.0001 |
| 4C | Drd1-Cre$^{+/-}$ mTOR$^{\Delta/Wt}$ Rosa26dTomato$^{+/-}$ (n=9)<br>Drd1-Cre$^{+/-}$ mTOR$^{\Delta/LoxP}$ Rosa26dTomato$^{+/-}$ (n=9) | Kruskal-Wallis test; p=0.0003 |
| 4D | Drd1-Cre$^{+/-}$ mTOR$^{\Delta/Wt}$ Rosa26dTomato$^{+/-}$ (n=9)<br>Drd1-Cre$^{+/-}$ mTOR$^{\Delta/LoxP}$ Rosa26dTomato$^{+/-}$ (n=9) | Kruskal-Wallis test; p<0.0001 |
| 4E | Drd1-Cre$^{+/-}$ mTOR$^{\Delta/Wt}$ Rosa26dTomato$^{+/-}$ (n=9)<br>Drd1-Cre$^{+/-}$ mTOR$^{\Delta/LoxP}$ Rosa26dTomato$^{+/-}$ (n=9) | Kruskal-Wallis test; p=0.7557 |
| 4F | Drd1-Cre$^{+/-}$ mTOR$^{\Delta/Wt}$ Rosa26dTomato$^{+/-}$ (n=9)<br>Drd1-Cre$^{+/-}$ mTOR$^{\Delta/LoxP}$ Rosa26dTomato$^{+/-}$ (n=9) | Kruskal-Wallis test; p=0.0052 |
| 5A | Drd1-Cre$^{+/-}$ mTOR$^{\Delta/Wt}$ Rosa26dTomato$^{+/-}$ (n=9)<br>Drd1-Cre$^{+/-}$ mTOR$^{\Delta/LoxP}$ Rosa26dTomato$^{+/-}$ (n=9) | Kruskal-Wallis test; p=0.7127 |
| 5B | Drd1-Cre$^{+/-}$ mTOR$^{\Delta/Wt}$ Rosa26dTomato$^{+/-}$ (n=9)<br>Drd1-Cre$^{+/-}$ mTOR$^{\Delta/LoxP}$ Rosa26dTomato$^{+/-}$ (n=9) | Kruskal-Wallis test; p<0.0001 |
| 5C | Drd1-Cre$^{+/-}$ mTOR$^{\Delta/Wt}$ Rosa26dTomato$^{+/-}$ (n=9)<br>Drd1-Cre$^{+/-}$ mTOR$^{\Delta/LoxP}$ Rosa26dTomato$^{+/-}$ (n=9) | Kruskal-Wallis test; p<0.0001 |
| 5D | Drd1-Cre$^{+/-}$ mTOR$^{\Delta/Wt}$ Rosa26dTomato$^{+/-}$ (n=9)<br>Drd1-Cre$^{+/-}$ mTOR$^{\Delta/LoxP}$ Rosa26dTomato$^{+/-}$ (n=9) | Kruskal-Wallis test; p<0.0001 |

| | | |
|---|---|---|
| 5E | Drd1-Cre$^{+/-}$ mTOR$^{\Delta/Wt}$ Rosa26dTomato$^{+/-}$ (n=9)<br>Drd1-Cre$^{+/-}$ mTOR$^{\Delta/LoxP}$ Rosa26dTomato$^{+/-}$ (n=9) | Kruskal-Wallis test; p=0.8493 |
| 5F | Drd1-Cre$^{+/-}$ mTOR$^{\Delta/Wt}$ Rosa26dTomato$^{+/-}$ (n=9)<br>Drd1-Cre$^{+/-}$ mTOR$^{\Delta/LoxP}$ Rosa26dTomato$^{+/-}$ (n=9) | Kruskal-Wallis test; p<0.0001 |
